# ModelTest-NG: a new and scalable tool for the selection of DNA and protein evolutionary models

**DOI:** 10.1101/612903

**Authors:** Diego Darriba, David. Posada, Alexey M. Kozlov, Alexandros Stamatakis, Benoit Morel, Tomas Flouri

## Abstract

ModelTest-NG is a re-implementation from scratch of jModelTest and ProtTest, two popular tools for selecting the best-fit nucleotide and amino acid substitution models, respectively. ModelTest-NG is one to two orders of magnitude faster than jModelTest and ProtTest but equally accurate, and introduces several new features, such as ascertainment bias correction, mixture and FreeRate models, or the automatic processing of partitioned datasets. ModelTest-NG is available under a GNU GPL3 license at https://github.com/ddarriba/modeltest.

---

It is well known that the use of distinct probabilistic models of evolution can change the outcome of phylogenetic analyses (Buckley, 2002; Buckley and Cunningham, 2002; Lemmon and Moriarty, 2004). Not surprisingly, a number of bioinformatic tools have been developed in the last 20 years for selecting the best-fit model for the data at hand (Darriba *et al.*, 2011, 2012; Kalyaanamoorthy *et al.*, 2017). Recently, Abadi *et al.* (2019) concluded that using a parameter-rich model for DNA data leads to very similar inferences as the best-fit models. The authors present the results as an average over a number of MSAs. However, looking at individual MSA analyses we may observe substantial topological distances between inferences under the best-fit models and inferences under a parameter-rich GTR model (Arbiza *et al.*, 2011; Hoff *et al.*, 2016). However, continuous advances in sequencing technologies have made possible the assemblage of large multiple sequence alignments (MSA) that require faster and more scalable tools. In particular, our tools ProtTest (Darriba *et al.*, 2011) and jModelTest (Darriba *et al.*, 2012), which are among the most popular tools for DNA and protein model selection, despite implementing High Performance Computing (HPC) algorithms for parallel execution with dynamic load balancing, still rely on PhyML (Guindon and Gascuel, 2003) for calculating the maximum likelihood (ML) scores of the competing models. This step constitutes by far the most compute-intensive part, >99% of overall execution time. The dependency on PhyML has several disadvantages: First, software maintenance heavily depends on PhyML developers. Thus, bug fixes and modifications affecting the interplay between Prot/jModelTest and PhyML must be continuously taken into account when incorporating PhyML updates. Second, PhyML and hence, ProtTest and jModelTest are relatively inefficient with respect to their likelihood calculations compared to more recent tools such as IQ-TREE (Nguyen *et al.*, 2015). The model selection feature of IQ-TREE, called ModelFinder (Kalyaanamoorthy *et al.*, 2017) is becoming increasingly popular due to its algorithmic and computational efficiency, the wide range of supported evolutionary models, and its user-friendliness.

With all this in mind, here we introduce ModelTest-NG, a new program that outperforms existing model selection tools in terms of the speed/accuracy trade-off. ModelTest-NG offers a completely redesigned graphical user interface (GUI). Recently, we also integrated ModelTest-NG into ParGenes (Morel *et al.*, 2018), a pipeline for massively parallel gene model selection and gene tree inference which is computationally efficient as well as easy-to-use. Moreover, we significantly improved maintainability, user support, and added several new capabilities. Its main features are:

- Data and models supported: ModelTest-NG supports both, nucleotide and amino acid models. It uses statistical criteria for selecting the best-fit substitution models such as AIC (Akaike, 1974), BIC (Schwarz, 1978), and DT (Minin *et al.*, 2003). For DNA, it can select among all 203 possible time-reversible substitution models. For amino acid data, ModelTest-NG compares 19 empirical replacement matrices as well as several recently introduced mixture models such as LG4M and LG4X (Le *et al.*, 2012). ModelTest-NG can also assess the fit of FreeRate models (Soubrier *et al.*, 2012).
- Partitioned data sets: ModelTest-NG can automatically perform model selection on individual non-overlapping partitions as specified by the user (e.g., on a per-gene basis, or by codon position).
- Phylogenetic templates: users can select so-called templates for the most popular phylogenetic inference tools: RAxML (Stamatakis, 2014), RAxML-NG (Kozlov *et al.*, 2018), IQ-TREE, PhyML, PAUP (Swofford, 2002), MrBayes (Ronquist *et al.*, 2012). When such a template is specified, ModelTest-NG will only evaluate models supported by the corresponding tool, and print out the respective tool-specific command line for phylogenetic reconstruction under the best-fit model.
- Native implementation: ModelTest-NG constitutes a full re-implementation of jModelTest and ProtTest in C++ that relies on a novel and efficient low-level implementation of the Phylogenetic Likelihood Library (https://github.com/xflouris/libpll-2). This library encapsulates all compute- and memory-intensive phylogenetic likelihood computations and fully leverages the capabilities of modern x86 processors by using the AVX and AVX2 vector instruction sets. It also incorporates a recent algorithmic technique for accelerating likelihood calculations (Kobert *et al.*, 2017). All required numerical optimization routines are implemented in the pll-modules library (https://github.com/ddarriba/pll-modules).

We benchmarked ModelTest-NG against jModelTest, ProtTest, and ModelFinder (part of IQ-TREE version 1.6.1) using empirical as well as simulated data sets. We measured runtimes for all datasets, and estimated accuracy (recovering the generating model) using the simulated datasets. The data sets and the experimental setup are described in detail in the supplementary material where we also discuss the results more extensively. For the empirical DNA data sets, ModelTest-NG yielded average speedups of 281.54 over jModelTest and of 1.12 over ModelFinder (Figure S1). For the simulated DNA data, ModelTest-NG was 110.77 times faster than jModelTest but slower than ModelFinder (the latter was 1.59 times faster than ModelTest-NG). On the empirical protein data sets, ModelTest-NG yielded average speedups of 36.94 over ProtTest and similar runtimes as ModelFinder. On the simulated protein data, ModelTest-NG was 36.07 times faster than ProtTest and 1.03 times faster than ModelFinder. We observed that ModelTest-NG scales better than ModelFinder and jModelTest/ProtTest on large data sets. In general, the larger the data set is, in terms of number of taxa and number of sites, the better ModelTest-NG performs compared to the competing tools (see Figure 1). In terms of accuracy, ModelTest-NG found the true generating model for 81% of the simulated DNA data sets (jModelTest: 81%, ModelFinder: 70%) and for 85% of the simulated protein data sets (ProtTest: 85%, ModelFinder: 87%) (Figure 1).

**FIG. 1.**
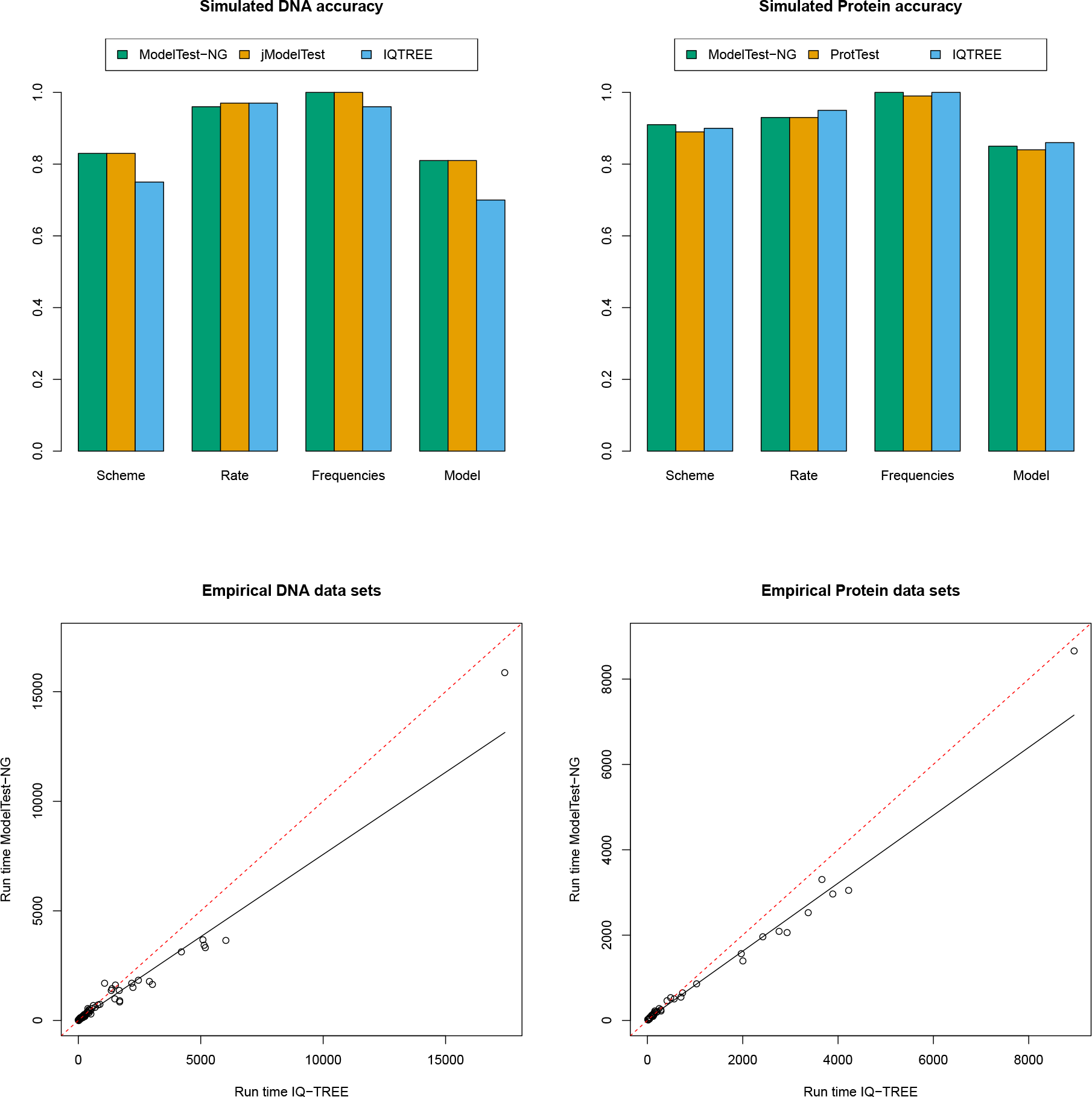
Model selection accuracy comparison between ModelTest-NG, jModelTest/ProtTest, and IQ-TREE for simulated data (top) and LOESS curved fitted to a scatter plot of ModelTest-NG run times versus IQ-TREE for empirical data (bottom), for DNA (left) and protein (right) MSAs. The dashed red line represents equal run times.

These results were obtained under the default model selection parameter settings for both tools. In additional experiments we found that there is a pronounced trade-off between speed and accuracy. Thus, we can expect that the more thoroughly we optimize the likelihood score for a set of substitution model parameters, the more accurate the results will become (see Supplementary Material online). The thoroughness of model parameter optimization routines can be specified in ModelTest-NG.

ModelTest-NG thus represents a substantial improvement over our previous highly-cited tools, jModelTest and ProtTest. It preserves the accuracy of its predecessors, as evaluated against the ground truth on simulated datasets, while the runtime is improved by two orders of magnitude on empirical data. Compared to IQ-TREE we observed similar run times for empirical data sets, but IQ-TREE was faster on synthetic data and particularly so on DNA data. However, the accuracy of IQ-TREE on DNA data was substantially lower than for ModelTest-NG (70% versus 81%, respectively). In future versions of ModelTest-NG, we intend to introduce methods to dynamically determine the optimal speed/accuracy trade-off for the dataset at hand.

## Supporting information

ModelTest-NG Supplementary Material

## Supplementary Material

Supplementary tables S1–S5 and figures S1–S4 are available at BIORXIV.

## Acknowledgments

This work was supported by the Ministry of Economy and Competitiveness of Spain and FEDER funds of the EU (Project TIN2016-75845-P) and by the Galician Government (Xunta de Galicia) under the Consolidation Program of Competitive Research (ref. ED431C 2017/04). Part of this work was funded by the Klaus Tschira foundation and DFG grant STA-860/6.

