## Supplementary material for "ModelTest-NG: a new and scalable tool for the selection of DNA and protein evolutionary models": ModelTest-NG Supplementary Material

#### Supplementary Data

#### 1. Test data sets

For the runtime and accuracy assessments, we used five different collections empirical and simulated DNA and protein multiple sequence alignments (Table S1).

1. **Empirical RAxML DNA**. This is the empirical DNA data previously used for benchmarking RAxML (Stamatakis 2014) (available at https://github.com/stamatak/test-Datasets).
2. **Empirical IQTREE DNA.** This is the empirical DNA data used for benchmarking IQ-TREE (available at http://www.iqtree.org/ModelFinder/).
3. **Empirical IQTREE Protein.** This is the empirical protein data used for benchmarking IQ-TREE (available at http://www.iqtree.org/ModelFinder/).
4. **Simulated DNA.** This is the simulated DNA data generated in Darriba et al., 2012.
5. **Simulated Protein.** This is a simulated protein data set generated following the process described in Darriba et al., 2012.

The properties of these test data set collections are summarized in Table S1.

*Table S1: Summary of test data set collections. `Count` denotes the number of MSAs, `Taxa` is the number of sequences, `Sites` is the number of columns, and `Patterns` is the number of unique site patterns. Note that the actual number of unique site patterns determines the computational requirements, as identical sites are typically compressed into site patterns by all common phylogenetic inference tools.*

| | | Count | Taxa | | | Sites | | | Patterns | | |
|---|---|---|---|---|---|---|---|---|---|---|---|
| | | | Min | Avg | Max | Min | Avg | Max | Min | Avg | Max |
| DNA | Empirical RAxML | 31 | 59 | 668 | 3782 | 297 | 2826 | 29198 | 234 | 1958 | 19437 |
| | Empirical IQTREE | 50 | 42 | 253 | 699 | 485 | 8428 | 93789 | 138 | 3129 | 32724 |
| | Simulated | 1000 | 10 | 55 | 100 | 501 | 999 | 1497 | 129 | 724 | 1491 |
| PROT | Empirical IQTREE | 45 | 26 | 81 | 194 | 127 | 2830 | 21155 | 108 | 2533 | 15022 |
| | Simulated | 1000 | 10 | 54 | 100 | 100 | 654 | 1200 | 46 | 481 | 1185 |

### 2. Evaluation setup

We compared ModelTest-NG version 0.1.5 with jModelTest version 2.1.10, ProtTest version 3.4.2 and IQTREE version 1.6.1 (note that the dedicated model selection module in IQTREE is called ModelFinder). For the sake of simplicity, all programs were executed in sequential execution mode using a single thread on a single physical core.

For DNA models, we used the GTR family of 88 nested models described in *Posada 2008*: 11 substitution schemes (JC/F81, K80/HKY, TrN, TPM1, TPM2, TPM3, TIM1, TIM2, TIM3, TVM, SYM/GTR) times four configurations of among-site rate variation (uniform, +I, +G, +I+G) times 2 stationary frequency models (equal or maximum-likelihood estimate). The command lines used were the following:

- modeltest-ng -i <msa_file> -t mp -d nt
- iqtree -s <msa_file> -m TESTONLY -mfreq FO
- jModelTest.jar -d <msa_file> -t BIONJ -S 11 -i -f -g4 -BIC -AIC -AICc -DT -tr 1

For protein models, we compared the 144 models available by default in ProtTest and in IQ-TREE version 1.6.1: 18 empirical substitution matrices times four configurations of among-site rate variation (uniform, +I, +G, +I+G) times two stationary frequencies (fixed by the model or empirical from the input data). The STMTREV model was excluded from the ModelTest-NG analysis as it was neither included in IQ-TREE nor in ProtTest. For real data sets, we also included LG4X and LG4M mixture models. The command lines we used were the following:

- modeltest-ng -i <msa_file> -t mp -d aa -m -STMTREV,+LG4M,+LG4X
- iqtree -s <msa_file> -st AA -m TESTONLY -madd LG4X,LG4M
- ProtTest.jar -i <msa_file> -I -G -IG -F -BIC -AIC -AICC -DT -S 0 -threads 1"

To evaluate performance we measured the run times. The execution times for ModelTest-NG, jModelTest, and ProtTest correspond to the overall wall clock time. Note that IQTREE also optimizes model parameters for the selected best-fit model after the model selection process. Therefore, to conduct a fair comparison, we only measured the time required for the actual model selection process.

We measure speedups via two distinct metrics: “local average” is the average speedup over all individual per-MSA speedups, and “global” is the overall speedup for analyzing all MSAs in the respective test data set (i.e., the ratio of accumulated runtimes for evaluating the entire test data set). This global speedup value better reflects the potential computational savings since a speedup of say 1.2 will yield substantially larger CPU resource savings on a large MSA that takes hours to analyze than on a small MSA that requires a few seconds to run.

For each test data set collection, we also compared the respective model selection results (best-fit model, substitution scheme, among-site rate variation model, stationary frequencies).

### 3. Computing infrastructure used

We executed all benchmarks on an Intel i7-2600 system with 4 physical cores, equipped with 16GB RAM and swapping disabled (see Table S2 for details).

*Table S2: Technical data of the computing cluster used for our experiments.*

| Hardware | | Software | |
|---|---|---|---|
| CPU model | Intel i7-2600 | OS | Ubuntu 16.04.5 LTS |
| CPU Architecture | Sandy Bridge | | |
| Cores | 4 @ 3.40GHz | Compiler | Gcc 5.2.0 |
| Memory | 16GB DDR3 | | |
| Vector Extensions | AVX, SSE4 | | |

### 4. Results

4.1 Speed up

Table S3 below shows the speedups between ModelTest-NG and the competing tools for all test data sets.

Our evaluations show that ModelTest-NG is on average 1 to 2 orders of magnitude faster than both jModelTest, and ProtTest. In particular, for large MSAs in terms of the number of sequences/taxa (e.g., MSAs included in the DNA RAxML test data set collection), differences in runtimes were even more pronounced. For some MSAs in the **Empirical RAxML DNA** test data set collection, we observed speedups exceeding a factor of 3,000.

Figure S1 shows the runtimes. Compared to IQTREE, ModelTest-NG exhibited slightly better run times for the empirical DNA test data sets (average speedup of 1.16 and 1.08 for the RAxML and IQTREE test data sets, respectively). For the empirical protein test data set, the average speedup was 1.00. The global speedup was generally better than the local average speedup, indicating that ModelTest-NG scales better than IQ-Tree on larger data sets (Figure S2).

*Table S3: Local and global speedups between ModelTest-NG and competing tools. "Local Average" is the average speedup over all individual per-MSA speedups, and "Global" is the overall speedup for analyzing all MSAs in the respective test data set (i.e., ratio of accumulated runtimes for evaluating the entire test data set)*

| | ProtTest/jModelTest | | IQTREE | |
|---|---|---|---|---|
| | Local Average | Global | Local Average | Global |
| Simulated DNA | 116.23 | 110.77 | 0.63 | 0.63 |
| Empirical DNA RAxML | 369.83 | 855.39 | 1.16 | 1.25 |
| Empirical DNA IQTREE | 193.23 | 164.87 | 1.08 | 1.24 |
| Simulated Protein | 36.79 | 36.07 | 0.47 | 1.03 |
| Empirical Protein IQTREE | 36.94 | 42.31 | 1.00 | 1.19 |

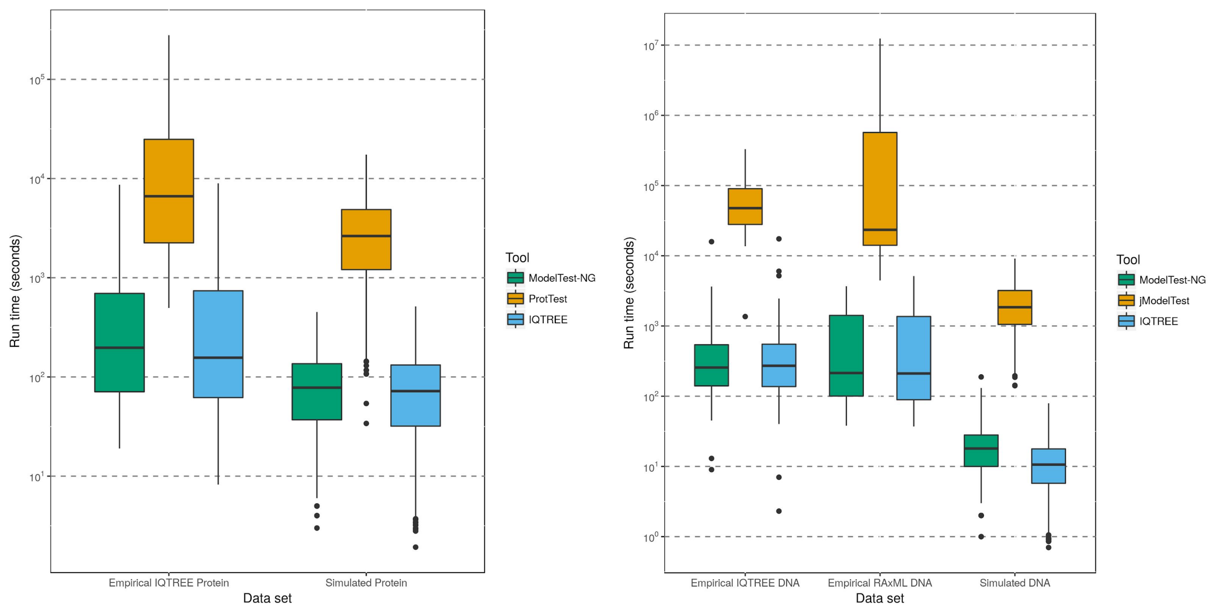


*Figure S1: Run time comparison between ModelTest-NG, jModelTest/ProtTest, and IQ-TREE, for empirical and simulated DNA (left) and protein (right) MSAs.Note the logarithmic scale on the 'y' axis.*

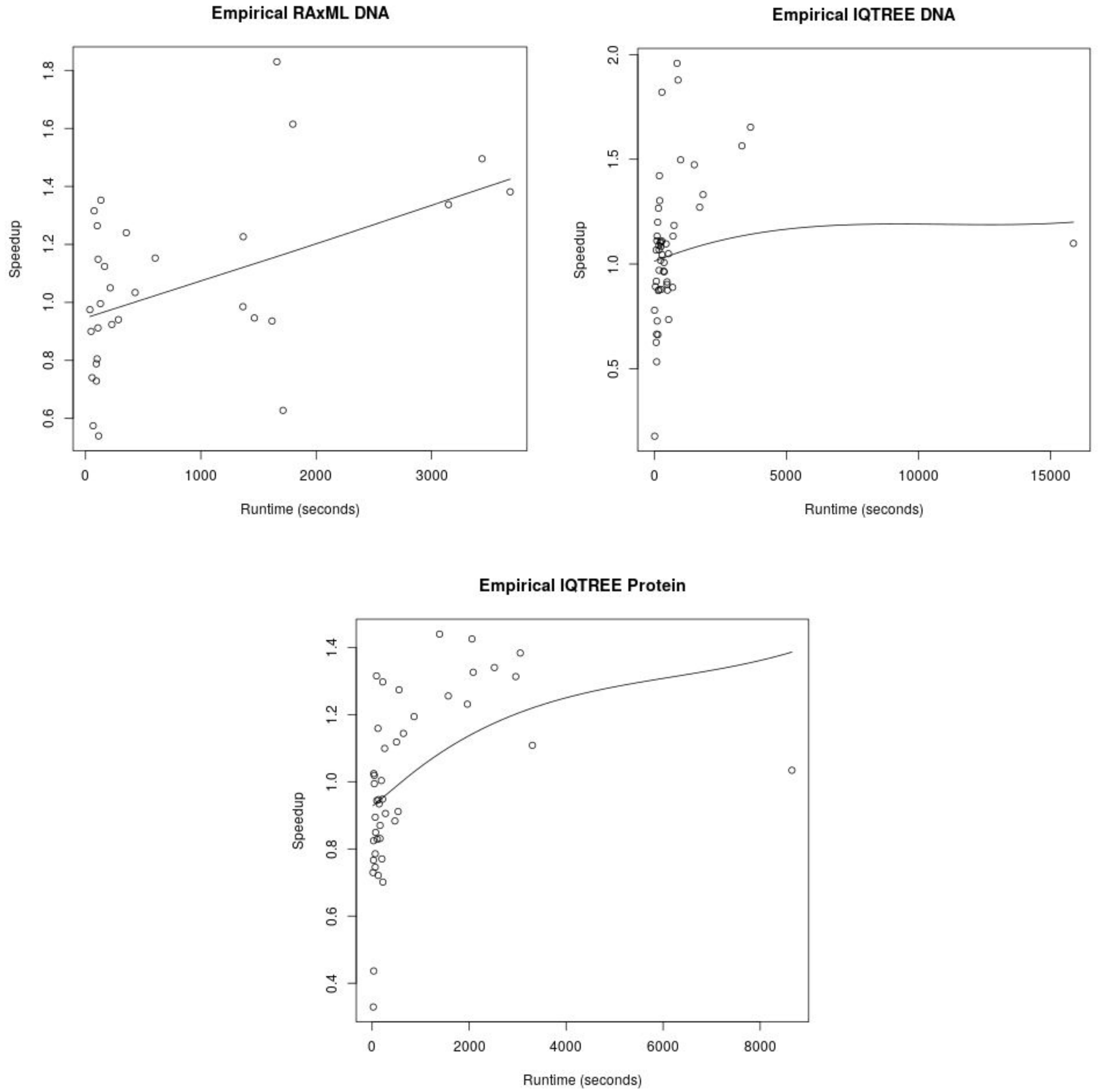


*Figure S2: LOESS curve fitted to a scatter plot of ModelTest-NG speedups versus IQTREE according to the ModelTest-NG runtimes for empirical data sets.*

4.2 *Accuracy*

In our comparison of the selected best-fit models on simulated data against the known ground truth (Table S4 and Figure S3), ModelTest-NG shows analogous results to jModelTest and ProtTest, identifying the true generating model for 81% and 84% of  the simulated DNA and protein MSAs, respectively. IQTREE performed worse (70%), while for protein data its performance was analogous to ModelTest-NG as well as ProtTest (also 84%).

If we compare the results obtained by the different tools for the RAxML test data set collection (Table S4), we observe that ModelTest-NG and jModelTest attain 100% agreement  with respect to the selected substitution scheme and among site rate variation model. ModelTest-NG and IQTREE agree on the best-fit model for 90% of the MSAs.

Overall, ModelTest-NG is slightly faster than IQ-Tree on empirical datasets, whereas on simulated data it is slightly slower. However, for simulated DNA data, the increased speed of IQ-Tree comes at the cost of a slight decrease in accuracy. The general trend is that ModelTest-NG shows higher speedups with increasing MSA size, both in terms of number of taxa and MSA patterns.

*Table S4. Model selection results comparison against the `ground truth` (for simulated data sets) and against ModelTest-NG. Values show the fraction by which the models selected by the tools agree. `Model` is the overall substitution model (i.e., combination of `Scheme`, `Rate` and `Freqs`). `Scheme` is the substitution scheme (DNA) or empirical substitution matrix (protein data). `Rate` is the among site rate variation model . `Freqs` is the stationary frequency type (equal or ML estimate for DNA data sets; defined by the model or empirical for protein data sets)*

| | | | Accuracy | | | | | | | |
|---|---|---|---|---|---|---|---|---|---|---|
| | | | Vs Ground Truth | | | | MTNG vs … | | | |
| | | | Model | Scheme | Rate | Freqs | Model | Scheme | Rate | Freqs |
| DNA | SIM | MTNG | 0.81 | 0.83 | 0.96 | 1.00 | - | - | - | - |
| | | jMT | 0.81 | 0.84 | 0.97 | 1.00 | 0.97 | 0.99 | 0.98 | 1.00 |
| | | IQTREE | 0.70 | 0.75 | 0.97 | 0.97 | 0.83 | 0.89 | 0.98 | 0.96 |
| | RML | MTNG | - | - | - | - | - | - | - | - |
| | | jMT | - | - | - | - | 0.87 | 1.00 | 1.00 | 0.87 |
| | | IQTREE | - | - | - | - | 0.90 | 0.94 | 1.00 | 0.97 |
| | IQT | MTNG | - | - | - | - | - | - | - | - |
| | | jMT | - | - | - | - | 0.91 | 0.94 | 1.00 | 0.94 |
| | | IQTREE | - | - | - | - | 0.94 | 0.94 | 1.00 | 1.00 |
| AA | SIM | MTNG | 0.85 | 0.90 | 0.93 | 1.00 | - | - | - | - |
| | | PT | 0.85 | 0.90 | 0.93 | 0.99 | 0.94 | 0.97 | 0.96 | 0.99 |
| | | IQTREE | 0.87 | 0.90 | 0.95 | 1.00 | 0.97 | 0.99 | 0.98 | 1.00 |
| | IQT | MTNG | - | - | - | - | - | - | - | - |
| | | PT | - | - | - | - | 0.91 | 0.98 | 0.93 | 1.00 |
| | | IQTREE | - | - | - | - | 0.89 | 0.93 | 0.96 | 1.00 |

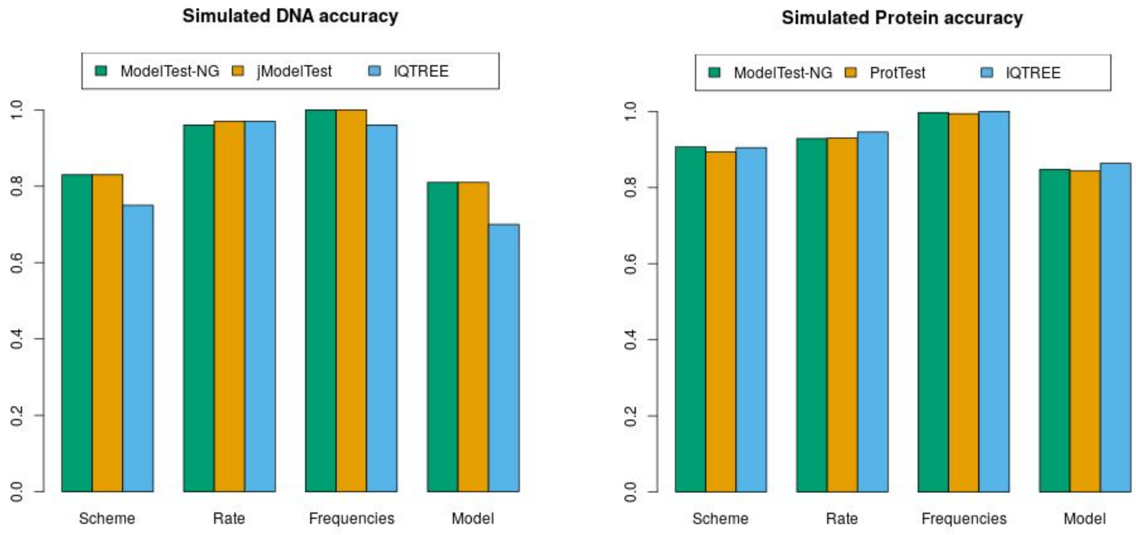


*Figure S3. Model selection accuracy results. Comparison against the `ground truth` (for simulated data sets). "Scheme" is the substitution scheme (i.e., rate matrix symmetries for DNA data and empirical matrix for protein data), "Rate" is the configurations of among-site rate variation, "Frequencies" is the stationary frequencies model, and "Model" is the combination of the previous 3 parameters.*

*Model Optimization Thoroughness*. ModelTest-NG optimizes model parameters with a tolerance of 0.01 log-Likelihood units by default. This tolerance provides a 'good' trade-off between selection accuracy and speed. To further investigate the impact of this value on execution times, we re-analyzed the simulated DNA MSAs using distinct tolerance settings. We used tolerances of 0.01, 0.1, and 0.5 log-likelihood units, and the observed model inference accuracy was 81%, 76%, and 66% respectively. The model inference accuracy was calculated as the fraction of samples where the true generating model was found. We observe that, if we increase this tolerance setting, both, execution time, and accuracy decreases (Table S5 and Figure S3).

*Table S5. Model selection results comparison against the `ground truth` (for simulated data sets) for ModelTest-NG, according to the model parameter optimization tolerance (in log-Likelihood units). Values show the fraction by which the models selected by the tools agree. `Model` is the overall substitution model (i.e., combination of `Scheme`, `Rate` and `Freqs`). `Scheme` is the substitution scheme (DNA) or empirical substitution matrix (protein data). `Rate` is the among site rate variation model . `Freqs` is the stationary frequency type (equal or ML estimate for DNA data sets; defined by the model or empirical for protein data sets). `Runtime` is the average runtime among the MSAs in the data set.*

| Tolerance | Model | Scheme | Rate | Freqs | Runtime |
|---|---|---|---|---|---|
| **0.01** | 0.805 | 0.834 | 0.961 | 1.000 | 20.690 |
| **0.10** | 0.764 | 0.821 | 0.925 | 1.000 | 20.140 |
| **0.50** | 0.660 | 0.751 | 0.880 | 0.995 | 16.890 |

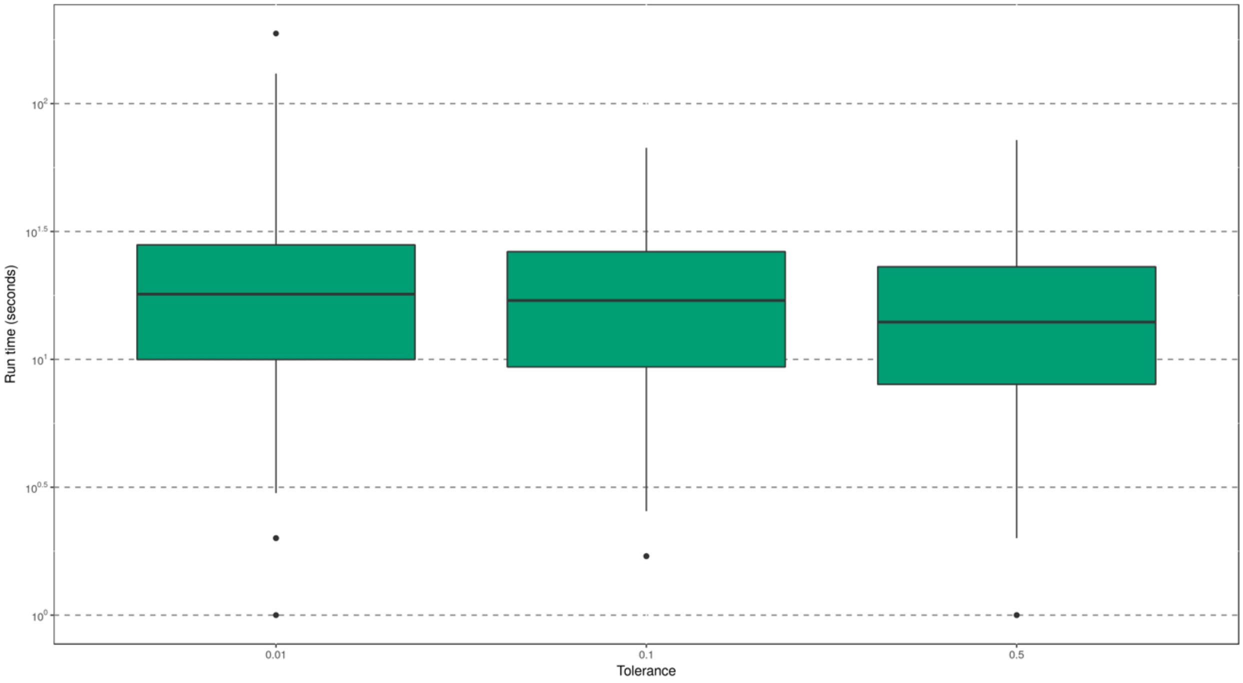


*Figure S4. ModelTest-NG run times according to the model optimization tolerance (in log-Likelihood units). Note the logarithmic scale on the y-axis.*
